# VIP interneurons selectively enhance weak but behaviorally-relevant stimuli

**DOI:** 10.1101/858001

**Authors:** Daniel J. Millman, Gabriel Koch Ocker, Shiella Caldejon, India Kato, Josh D. Larkin, Eric Kenji Lee, Jennifer Luviano, Chelsea Nayan, Thuyanh V. Nguyen, Kat North, Sam Seid, Cassandra White, Jerome A. Lecoq, R. Clay Reid, Michael A. Buice, Saskia E.J. de Vries

## Abstract

Vasoactive intestinal peptide-expressing (VIP) interneurons in cortex regulate feedback inhibition of pyramidal neurons through suppression of somatostatin-expressing (SST) interneurons and, reciprocally, SST neurons inhibit VIP neurons. Here, we show that VIP neurons in mouse primary visual cortex have complementary contrast tuning to SST neurons and respond synergistically to front-to-back visual motion and locomotion. Network modeling indicates that this VIP-SST mutual antagonism regulates the gain of cortex to achieve both sensitivity to behaviorally-relevant stimuli and network stability.

## Main Text

Inhibitory interneurons play a major role in establishing the dynamics of cortical microcircuits.^1,2^ In layer 2/3 of cortex, vasoactive intestinal peptide-expressing (VIP) interneurons regulate feedback inhibition of pyramidal neurons through suppression of somatostatin-expressing (SST) interneurons.^3^ Through this disinhibitory mechanism, VIP interneurons are believed to modulate network dynamics based on the behavioral state of the animal; for instance, VIP neurons in mouse primary visual cortex (V1) are reliably active during periods of locomotion.^4^ Moreover, VIP neurons in V1 are a target of top-down inputs and mediate enhancement of local pyramidal cell activity in response to activation of those inputs.^5^ VIP neurons also receive reciprocal inhibition from SST neurons, creating a circuit motif of mutual inhibition between VIP and SST neurons with unknown implications for cortical processing. Behaviorally, mouse V1 is necessary for the detection of low contrast visual stimuli,^6^ and the optogenetic activation of VIP neurons in mouse V1 increases contrast sensitivity whereas the activation of SST or PV neurons decreases it.^7^ This suggests that the perception of low contrast stimuli is strongly influenced by VIP neuron activity in V1. Although the activity of VIP neurons has been shown to be suppressed below baseline in response to high contrast full-field grating stimuli,^8^ the responses of VIP neurons to low contrast visual stimuli are not known. To this end, we investigated the influence of stimulus contrast and motor behavior (i.e. locomotion) on the visual responses of VIP, SST, and pyramidal neurons in mouse V1. SST neurons responded exclusively at high contrast whereas VIP neurons responded exclusively at low contrast with strong preference for front-to-back motion that is congruent with self-motion during locomotion. As a population, layer 2/3 – but not deeper layer – pyramidal neurons responded more strongly at low contrast than high contrast and showed a slight, but significant, bias for front-to-back motion. Finally, we made novel extensions of stabilized supralinear network (SSN) models to incorporate the diversity of inhibitory interneuron types and used these models to demonstrate that VIP-driven disinhibition at low contrast can drive large increases in pyramidal neuron activity, despite the relatively low activity of both SST and pyramidal neurons in this contrast regime. The selective enhancement of front-to-back motion could increase detection of obstacles approaching head-on during locomotion. Based on these results, we conclude that VIP neurons amplify responses of pyramidal neurons to weak but behaviorally-relevant stimuli.

We recorded responses to drifting gratings at eight directions and six contrasts during calcium imaging of mouse Cre lines for Vip and Sst as well as pyramidal neurons across cortical layers (Cux2: layer 2/3; Rorb: layer 4; Rbp4: layer 5; Ntsr1: layer 6) transgenically expressing GCaMP6f and computed the response to each stimulus condition. The majority of neurons were tuned for grating contrast and direction (bootstrapped χ^2^ test, p<0.01; Supplementary Figure 1a; Supplementary Table 1). We observed direction- or orientation-tuned neurons that responded preferentially either to high contrast gratings or low contrast gratings (Figure 1a). Substantial differences in contrast and direction tuning were apparent across Cre lines (Figure 1b-g). Virtually all VIP neurons responded only at low (<20%) contrast to front-to-back motion (0 degrees; nasal-to-temporal) or an adjacent direction, yielding the greatest direction bias among Cre lines as quantified by the vector sum of direction preferences (Figure 1c). The direction of bias was consistent across all VIP mice (n=6; Supplementary Figure 2a) and did not result from stimulus direction-selective running behavior (Supplementary Figure 2b). High contrast gratings of all directions suppressed VIP neuron activity (Figure 1e), consistent with a previous report^9^ of a small population of neurons that were active during locomotion but suppressed by the presentation of full-field high contrast gratings. SST neurons had high contrast selectivity, weak direction selectivity, and varied direction preference (Figure 1c,d,g), resulting in an average population response that was strong at high contrast across all directions, complementing the non-direction selective suppression at high contrast observed in VIP neurons. Unlike inhibitory interneurons, pyramidal neurons exhibited substantial direction and orientation selectivity and tiled all eight possible direction preferences (Figure 1b,f,g). Contrast preference among pyramidal neurons systematically varied across cortical layers, exhibiting a progression from a mixture of low and high contrast-preferring neurons in layer 2/3 to almost exclusively high contrast-preferring neurons in layers 5 and 6 (Figure 1b,d; Supplementary Figure 3). Like VIP neurons, CUX2 neurons in layer 2/3 showed direction bias toward front-to-back motion at 5% and 10% contrast but not higher contrasts (Figure 1c); pyramidal neurons in deeper layers did not have direction bias. Taken together, concerted changes in response magnitude near 20% contrast across all Cre lines and layers indicates the presence of a phase transition in cortical dynamics between a low contrast regime exemplified by relatively inactive SST neurons and a high contrast regime exemplified by highly active SST neurons.

**Figure 1:**
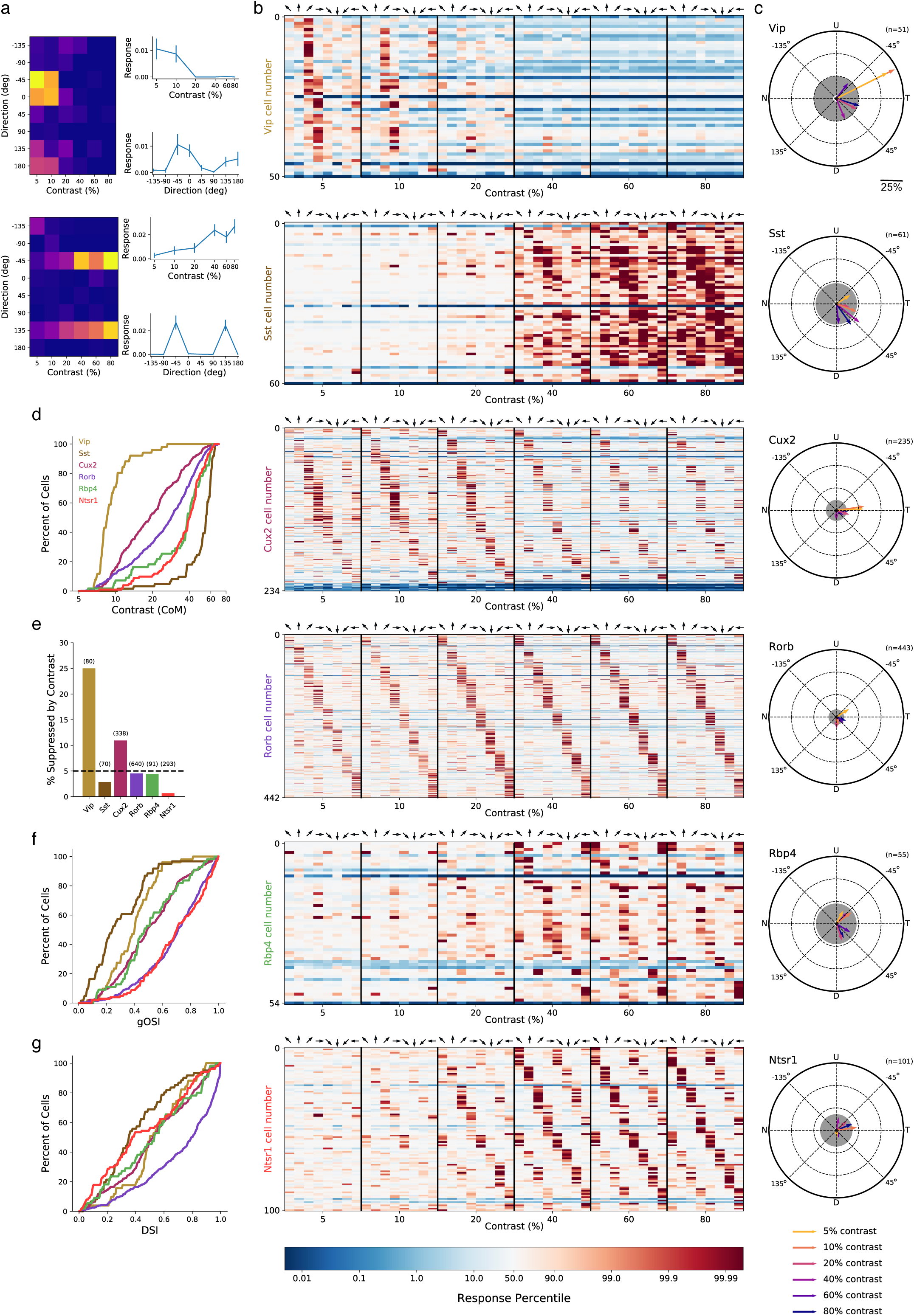
Contrast and direction preferences are cell-type and layer specific. **a)** Two examples of single cells, a low-pass (top) and a high-pass (bottom) contrast response function. Left: Heatmaps show the mean response of the cell averaged over all presentations of a drifting grating of a given direction and contrast. Right: Mean response to gratings of each contrast at the cell’s peak direction (top) as well as each direction at the cell’s peak contrast (bottom). Error bars are SEM. **b)** Waterfall plots showing the response significance at each contrast and direction of all responsive cells (χ^2^ test; p < 0.01) from mice of each Cre line. Cells are ordered by direction preference at the cell’s peak contrast. Response significance for each condition is obtained by comparing the mean condition response minus mean blank (i.e. zero contrast) response to a null distribution of such differences that is generated by shuffling responses across trials (see methods); responses below the median of the shuffle distribution are blue (i.e. suppressed), responses above the median of the shuffle distribution are red (i.e. enhanced). **c)** Radial plot of the average direction preference of cells of each Cre line at each contrast. Arrows are the vector sum of all responsive cells at a given contrast. Gray shaded region indicated a 90% confidence interval of the vector sum for population with uniformly-distributed direction preferences. Scale: The distance between each pair of concentric dashed rings is 25%. **d)** Cumulative distribution of contrast preferences (center-of-mass of a cell’s contrast response function; CoM) across Cre lines. **e)** Fraction of all cells of each Cre line that are suppressed by contrast. The mean response to all grating directions at 80% contrast must be significantly below mean blank response (bootstrapped distribution of mean response differences; family-wise type 1 error < 0.05; see methods). **f)** Cumulative distribution of global orientation selectivity indices (gOSI) across Cre lines. **g)** Cumulative distribution of direction selectivity indices across Cre lines.

To assess circuit-wide effects of locomotion on cortical dynamics, we examined the average activity of each neuron population as a whole. We focus here on the responses at low contrast in layers 2/3 and 4, but not layers 5 and 6 which did not respond at low contrast. VIP, SST, and pyramidal populations in both layers 2/3 and 4 all had increased activity during locomotion compared with stimulus presentations during which the mouse was stationary (Figure 2). During locomotion, the low contrast and front-to-back direction selectivity that was common to nearly all VIP neurons resulted in an average VIP population response that had tuning closely resembling the tuning of any individual VIP neuron (Figure 2a,e,i,m). By comparison, the VIP population did not respond to motion of any direction or contrast when the mice were stationary. Running also increased the SST population response to high contrast gratings, which also had the highest average response to front-to-back motion but responded strongly as a population to other directions as well (Figure 2b,f,j,n). The CUX2 population in layer 2/3 responded broadly across directions but more strongly, and with greater running enhancement, at low than high contrast (Figure 2c,g,k,o), whereas the Rorb population in layer 4 had comparable response magnitude and running enhancement across contrasts (Figure 2d,h,l,p).

**Figure 2:**
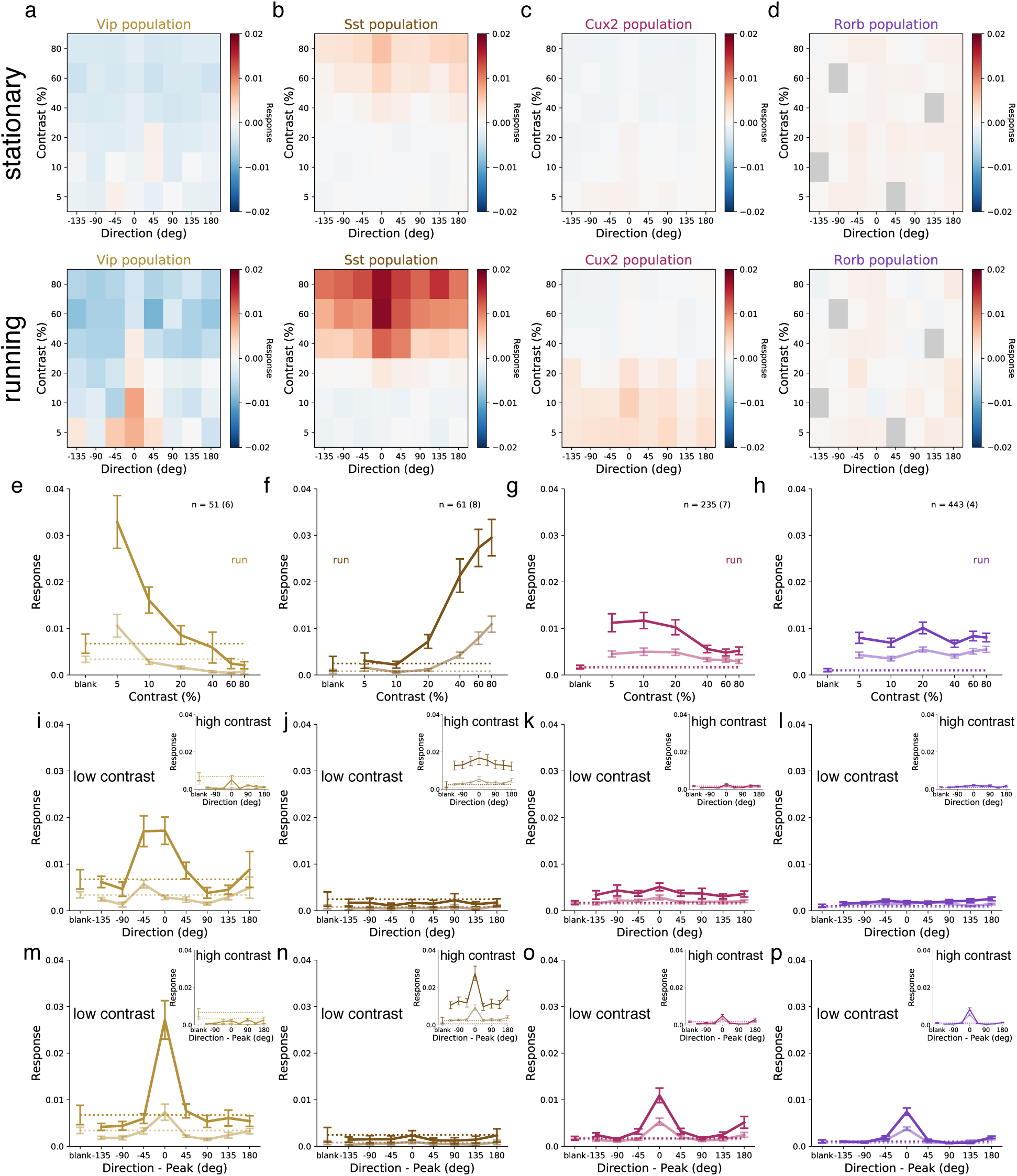
Average population responses of inhibitory, but not excitatory, cells are strongly biased toward front-to-back visual motion which is enhanced during locomotion. **a-d)** Mean blank-subtracted response magnitude of all cells from mice of each superficial Cre line during stationary (top) and running (bottom) periods. Gray boxes in Rorb plots indicate insufficient run and stationary data. **e-h)** Mean population contrast response tuning during stationary (faint lines) and running (bold lines) periods. **i-l)** Mean population direction response tuning at low (5-10%) contrast as in **e-h**. insets: mean population direction response tuning at high (60-80%) contrast. **m-p)** Mean single-cell direction tuning (i.e. aligned to each cell’s peak direction) as in **e-l**. All error bars are SEM. Sample size indicates number of cells with number of experiments in parenthesis.

Anatomical and optogenetic perturbation experiments suggest that VIP neurons disinhibit pyramidal neurons through inhibition of SST neurons.^3,5,10^ However, VIP neurons only respond to one direction of low contrast grating and SST neurons have very weak responses to low contrast gratings of any direction, potentially limiting the magnitude of SST activity that is available to be inhibited by VIP neurons and, consequently, limiting the magnitude of disinhibition of pyramidal neurons. Evidence that visual cortex has higher gain at low contrast than high contrast^11,12,13^ suggests that a small reduction in feedback inhibition (e.g. disinhibition) is capable to drive a large increase in pyramidal neuron activity. Stabilized supralinear network (SSN) models have been proposed to account for a variety of contrast-dependent response properties in visual cortex,^14,15^ including the transition from a high gain regime at low contrast to a feedback inhibition dominated low gain regime at high contrast.^16,17^ To investigate the distinct roles of each interneuron type, we extended the SSN model from one homogeneous population of interneurons to three populations corresponding to VIP, SST, and parvalbumin-expressing (PV) neurons to model layer 2/3 of mouse V1 (Figure 3a; Supplementary Methods). Briefly, the network is a ring model in which each CUX2 pyramidal neuron receives external (“sensory”) excitatory input that has Gaussian tuning with mean (i.e. peak/preferred direction) corresponding to the neuron’s position on the ring and standard deviation of 30 degrees; PV neurons also receive external input which is not tuned. The strength of external input is intended to represent a monotonically-increasing function of stimulus contrast, though no specific relationship is claimed here. Connections from CUX2 neurons (i.e. excitatory connections) also have Gaussian tuning that depends on the difference between the orientation preferences of the pre- and post-synaptic neurons whereas connections from inhibitory neurons (i.e. inhibitory connections) are not tuned (Figure 3b). All neurons are modeled as rate units with rectified quadratic transfer function. This model is able to qualitatively reproduce the population direction and contrast tuning we observed for VIP, SST, and CUX2 neurons as well as make a prediction for the tuning of PV neurons (Figure 3c). Model VIP neurons are most active at a low level of external input corresponding to the highest gain (“supralinear”) regime for CUX2 and PV activity (Figure 3d: left). Ablating the VIP-to-SST inhibitory connection, the only output of VIP neurons contained in the model, results in a large reduction in the gain and activity of VIP, CUX2, and PV populations at low input (Figure 3d: right). These results indicate that VIP disinhibition is capable of producing substantial increases in gain at low contrast despite low activity of the intermediate SST neuron population.

**Figure 3:**
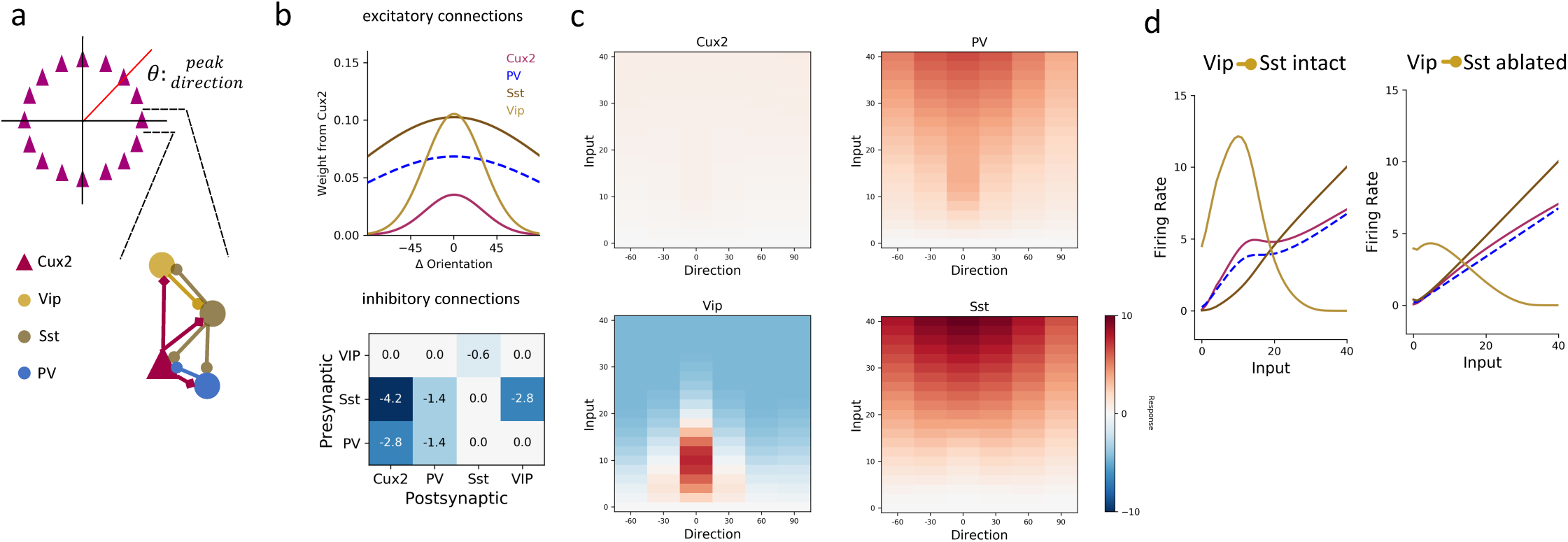
A stabilized supralinear network model with three interneuron populations reproduces contrast and direction tuning of multiple cell types and implicates Vip cells in enhancement of network gain for weak inputs. **a)** Top: The network architecture is a ring corresponding to the peak of each Cux2 pyramidal cell’s direction tuning curve. The entire ring spans 180 degrees of direction. Bottom: A schematic illustrates the connectivity among cell types. **b)** Top: The distribution of excitatory connection strength from Cux2 pyramidal cells onto each cell type is Gaussian with mean equal to the difference in orientation preference of pre- and post-synaptic cells. The distributions of recurrent connections onto Cux2 cells and connections onto Vip cells are narrow (standard deviation of 30 degrees) compared to the distributions onto PV and Sst cells (standard deviation of 100 degrees). Bottom: Inhibitory connection weights do not vary as a function of the difference between the peak directions of pre- and post-synaptic cells. **c)** The average population responses across direction and contrast conditions qualitatively reproduce experimental data for Cux2, Sst, and Vip cells shown in Figure 2. **d)** Left: The steady state firing rates are shown for model cells of each type with peak direction tuning of zero degrees in response to an input “stimulus” of zero degrees. Right: The steady state firing rates of the same model cells in response to an input of zero degrees with the Vip-to-Sst connection strength set to zero demonstrates that this connection is necessary for high gain of Cux2 and PV cells at the low input levels for which Vip cells are most responsive.

This survey of contrast tuning in mouse V1 revealed two distinct regimes of cortical dynamics in layer 2/3. At high contrast, SST neuron activity is high, VIP neuron activity is suppressed, and layer 2/3 pyramidal neuron activity is reduced compared with the low contrast regime; at low contrast, SST neuron activity is low, VIP neuron activity is direction tuned and gated by locomotion, and layer 2/3 pyramidal neuron activity is higher and more enhanced by locomotion. Measurements of size tuning have shown that SST neurons prefer large gratings, suggestive of a role mediating surround suppression, whereas VIP neurons only respond to gratings smaller than those that drive SST neurons.^18,19^ In mouse primary auditory cortex, VIP neurons are selective for lower sound intensities than SST or PV neurons.^20^ Taken together, a parsimonious explanation of these results is that VIP neuron activity supports a high gain regime that increases sensitivity to weak inputs, whereas SST neuron activity promotes a low gain regime that decreases sensitivity to strong inputs and maintains network stability. Heightened sensitivity to detect low contrast objects or obstacles approaching head-on during locomotion might be more behaviorally relevant than other directions of motion. This ability of VIP neurons to promote high gain in the local microcircuit might be indicative of a more general role at the nexus of top-down (e.g. attention) and bottom-up (e.g. saliency) processes.

## Author Contributions

Conceptualization, M.A.B. and S.E.J.dV. Methodology, D.J.M., G.K.O., J.A.L., and S.E.J.dV. Investigation, S.C, I.K., J.D.L., E.K.L., J.L., T.V.N., C.N., K.N., S.S., and C.W. Formal Analysis, D.J.M., G.K.O., and S.E.J.dV. Writing – Original Draft, D.J.M. Writing – Review & Editing, D.J.M., G.K.O., M.A.B., and S.E.J.dV. Supervision, J.A.L., R.C.R., M.A.B., and S.E.J.dV.

## Acknowledgements

The authors wish to thank the Allen Institute founder, Paul G. Allen, for his vision, encouragement and support.

---

**Supplementary Table 1:** The total number of cells and mice per Cre line.

|  | Cells (Mice) |
| --- | --- |
| Cux2 | 338 (7) |
| Rorb | 640 (4) |
| Rbp4 | 91 (4) |
| Ntsr1 | 293 (6) |
| Sst | 70 (8) |
| Vip | 80 (6) |

**Supplementary Figure 1:**
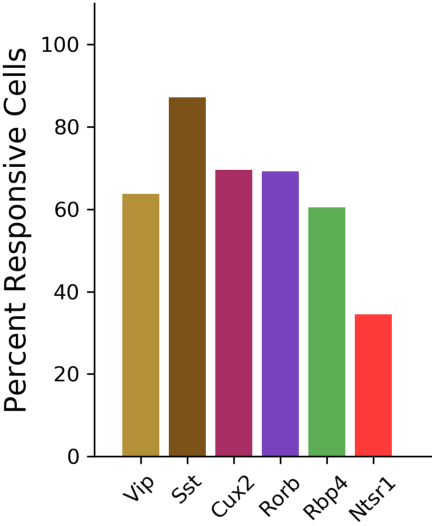
The fraction of imaged cells that were significantly responsive to the gratings stimulus (bootstrapped χ^2^ test, p<0.01).

**Supplementary Figure 2:**
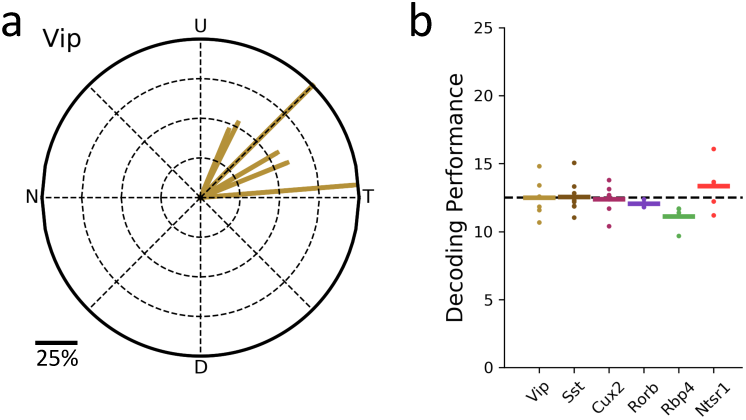
The direction of bias was consistent across all VIP mice and did not result from stimulus direction-selective running behavior. **a)** Vector sums for each of the six Vip-Cre mice. **b)** Performance of a linear support vector classifier trained to decode the direction of grating (1-of-8 classification) from the running speed of the mouse. The average validation performance for three-fold cross-validation is shown. Each dot is the performance for one mouse; bars are the mean across mice of a given Cre line.

**Supplementary Figure 3:**
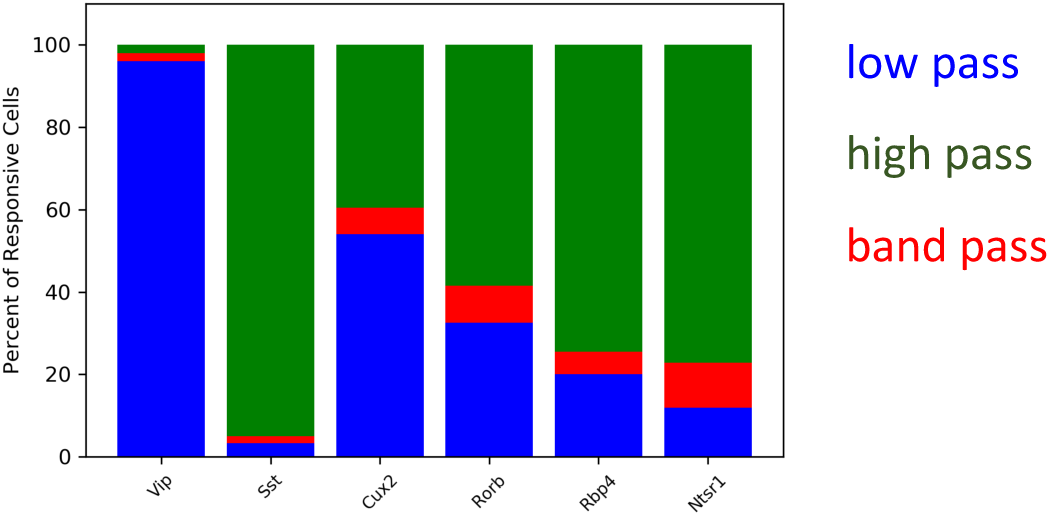
Distribution of contrast response types by Cre line determined by fitting of rising sigmoid (high pass), falling sigmoid (low pass), or the product of rising and falling sigmoids (band pass). See methods.

## Methods

### Experimental Animals

All animal procedures were approved by the Institutional Animal Care and Use Committee (IACUC) at the Allen Institute for Brain Science. Six double or triple transgenic mouse lines were used to drive expression of GCamp6/f in genetically-defined cell types, including four excitatory (Cux2-CreERT2;Camk2a-tTA;Ai93, Rorb-IRES2-Cre;Camk2a-tTA;Ai93, Rbp4-Cre_KL100;Camk2a-tTA;Ai93, and Ntsr1-Cre_GN220;Ai148) and two inhibitory (Vip-IRES-Cre;Ai148 and Sst-IRES-Cre;Ai148) mouse lines. Mice were habituated to head fixation and visual stimulus presentation for two weeks prior to data collection (See de Vries, Lecoq, Buice *et al.* for further Cre line, surgical, and habituation details).

### Two photon imaging platform and image processing

Data was collected using the same data collection pipeline as the Allen Brain Observatory and processed using the same image processing and event detection methods. All analyses of cell responses were performed on L0 penalized detected events (See de Vries, Lecoq, Buice *et al.* for further imaging and image processing details).

Briefly, two-photon imaging data was collected from the retinotopic center of primary visual cortex that was identified through mapping during widefield intrinsic signal imaging. Cux2-CreERT2;Camk2a-tTA;Ai93, Vip-IRES-Cre;Ai148, and Sst-IRES-Cre;Ai148 mice were imaged at 175 um below the cortical surface in layer 2/3; Rorb-IRES2-Cre;Camk2a-tTA;Ai93 mice were imaged at 275 um below the cortical surface in layer 4; Rbp4-Cre_KL100;Camk2a-tTA;Ai93 mice were imaged at 375 um below the cortical surface in layer 5; and Ntsr1-Cre_GN220;Ai148 mice were imaged at 550 um below the cortical surface in layer 6. (These Cre lines and imaging depths match those used in the Allen Brain Observatory.)

### Visual Stimulus

As experimental sessions took place on the same data collection pipeline as the Allen Brain Observatory, visual stimulus monitor calibration and positioning was identical (See de Vries, Lecoq, Buice *et al.* for further visual stimulus presentation details). The stimulus consisted of a full field drifting sinusoidal grating that was presented at a single spatial frequency (0.04 cycles/degree) and temporal frequency (1 Hz), 8 directions uniformly distributed in 45 degree increments (0 degrees = horizontal front-to-back motion), and 6 contrasts (5%, 10%, 20%, 40%, 60%, and 80%). Direction of motion was always orthogonal to the orientation of the grating. Each grating was presented for 2 seconds, followed by 1 second of mean luminance gray before the next grating. Each grating condition (direction, contrast combination) was presented 15 times. Trials were randomized with 30 randomly interleaved blank (i.e. mean luminance gray, zero contrast) trials.

## Analysis

### Statistical test for responsiveness

A chi-square test for independence was used to determine significantly responsive cells to the drifting grating stimulus set. A chi-square test statistic was computed 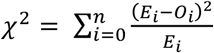, where 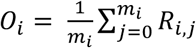 is the observed average response (*R*) of the neuron over *m* presentations of a grating stimulus of a particular condition (i.e. direction-by-contrast pair or blank, n = 49 total conditions), and 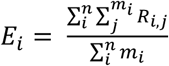 is the expected (grand average) response per stimulus presentation. A p-value was then calculated for each cell by comparing the test statistic against a null distribution of 200,000 test statistics, each computed from the cell’s responses after shuffling (with replacement) cell responses across all presentations.

### Response Significance by Stimulus Condition and test for suppression by contrast

The distribution of responses to stimulus presentations varied substantially across cells. To facilitate the visualization of responses across all cells and stimulus conditions simultaneously (figure 1b), a statistical measure was used to normalize response magnitudes. The mean blank-subtracted response to a given stimulus condition was calculated as: 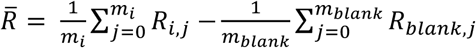. Then, a bootstrapped null distribution of such mean (blank-subtracted) condition responses was generated by sampling with replacement from all of the cell’s responses across all stimulus presentations. The percentiles of each cell’s observed mean condition response within its own bootstrapped distribution are the values plotted in Figure 1b. Cells were determined to be suppressed by high contrast if this percentile for the peak direction grating condition at 80% contrast was below 0.05.

### Orientation and Direction Selectivity Metrics

Global orientation selectivity was computed from mean extracted event responses to drifting gratings, at the cell’s preferred contrast, as

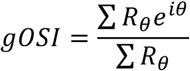

where *θ* is the direction of grating movement, and *R*_*θ*_ is the mean response to that direction of motion.

Direction selectivity was computed from mean extracted event responses to drifting gratings, at the cell’s preferred contrast, as

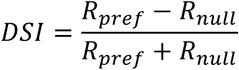

where *R*_*pref*_ is a cell’s mean response in its preferred direction (i.e. largest response-evoking direction) and *R*_*null*_ is its mean response to the opposite direction.

### Contrast Preference Metric

Contrast preference was computed from mean extracted event responses to drifting gratings, at the cell’s preferred direction, as

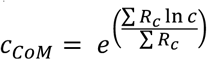

where c is the contrast of the drifting grating, *R*_*C*_ is a cell’s mean response at contrast c, and *C*_*COM*_ is the log-scaled center of mass of the cell’s contrast response tuning.

### Bias in population direction preference

The direction and magnitude of bias in direction preference for a population of cells (e.g. all cells recorded from one mouse or all cells recorded from all mice of a particular Cre line) was calculated as the direction and magnitude of the vector sum of the direction preferences of the cells that comprise the population, at a particular contrast, as

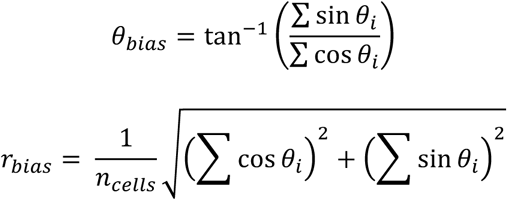

where *θ*_*i*_ is the preferred direction of cell *i*, *n*_*cells*_ is the number of cells in the population, *θ*_*bias*_ is the direction of the vector sum over the population, and *r*_*bias*_ is the magnitude of the vector sum over the population.

### Stimulus Tuning conditioned on locomotion behavior

As part of the standardized pipeline for the Allen Brain Observatory, mice were held on a running wheel during experimental sessions and locomotion behavior was recorded (See de Vries, Lecoq, Buice *et al.* for further run speed measurement details). The mean running speed was calculated for each trial over the same time window as the mean cellular response was calculated. Trials for which the mean running speed was greater than or equal to 1cm/s were categorized as running trials, whereas trials for which the mean running speed was below 1cm/s were categorized as stationary trials. The mean and standard error of the mean event magnitude for each contrast and direction condition shown in Figure 2 was calculated separately for running and stationary trials. The criterion for a cell to be included in the calculation for a given direction-by-contrast condition was that the mouse had to be running for a minimum of four trials *and* be stationary for a minimum of four trials of that condition.

### Contrast Response Function Fitting and model comparison

Event responses as a function of contrast, at a cell’s preferred direction, were fit to a rising sigmoid (“high pass”), a falling sigmoid (“low pass”), and the product of one rising and one falling sigmoid (“band pass”).

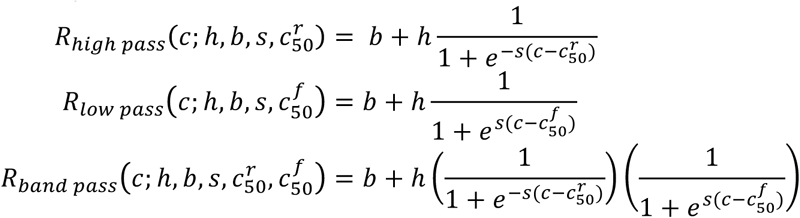

where *C* is the contrast, 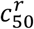 is the contrast at which the response rises halfway between the base and height, 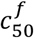 is the contrast at which the response falls halfway between the base and height, *b* is the lowest response, *h* is the response amplitude, and *s* is the slope of the sigmoid (fixed at *s* = 10). The best fit model was determined by calculating the Akaike Information Criterion (AIC) for each model and selecting the model with lowest AIC.

The AIC can be calculated as

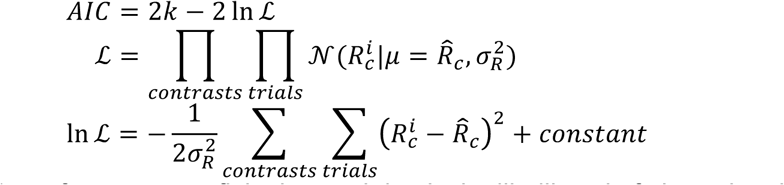

where *k* is the number of parameters fit in the model, ℒ is the likelihood of observing the responses given the fitted model and response distribution, 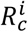 is the cell’s response to a grating stimulus of contrast c (at the cell’s preferred direction) on trial *i*, 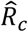 is the response predicted by the model to a grating stimulus of contrast c, 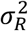 is the variance of all of the cell’s responses, and 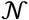 is the normal distribution. In practice, it is more convenient to directly calculate the log-likelihood than to calculate the likelihood and subsequently take the log, and the constant can be ignored for model selection since the same constant applies to all models being compared.

Due to the non-normal response distribution, possibly arising from calcium imaging as well as an underlying non-normal spiking distribution, we bootstrapped the log-likelihood rather than assume normality. Therefore, the likelihood was calculated numerically by shuffling responses across trials 1000 times and calculating the sum of square residuals from the predicted responses as 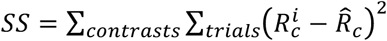 for each shuffle. The likelihood was taken as the fraction of shuffles for which *SS* was greater than the observed *SS*.

### Stabilized Supralinear Network (SSN) Model

The SSN was modelled as a ring network, largely maintaining the basic architecture and dynamics described in Rubin et al. (2015) but deviating primarily in the diversity of inhibitory neurons and distributions of connections between neuron populations (including untuned inhibitory connections, described below). Our network consisted of one excitatory population (representing layer 2/3 CUX2 pyramidal neurons) and three inhibitory populations (representing PV, SST, and VIP interneurons, respectively). The ring network structure was imposed by providing each excitatory neuron with external (“sensory”) excitatory input that had Gaussian tuning with mean (i.e. peak/preferred direction) corresponding to the neuron’s position on the ring and standard deviation of 30 degrees; PV neurons also received external input which was not tuned (i.e. all PV cells receive input of equal strength). The entire network covered 180 degrees of orientation (or direction). The strength of external input was intended to represent a monotonically-increasing function of stimulus contrast, though no specific relationship between input magnitude and contrast is claimed here.

Connections from CUX2 neurons (i.e. excitatory connections) also had Gaussian tuning that depended on the difference between the orientation preferences of the pre- and post-synaptic neurons, whereas connections from inhibitory neurons (i.e. inhibitory connections) were not tuned (Figure 3b). The distributions of recurrent connections onto Cux2 cells and connections onto Vip cells were narrow (standard deviation of 30 degrees) compared to the distributions onto PV and Sst cells (standard deviation of 100 degrees).

The network consisted of 184 excitatory neurons, 40 PV neurons, 15 SST neurons, and 15 VIP neurons. The excitatory population had 180 neurons with uniform 1-degree spacing of peak directions to tile the ring, plus 4 extra neurons with peak direction of zero degrees to capture the slight bias of the CUX2 neurons. All model VIP neurons had a peak direction of zero degrees to capture the strong bias for front-to-back motion observed for VIP neurons. In addition, all SST and PV model neurons also had a peak direction of zero degrees, though the very broadly-tuned inputs to these neurons results in a much weaker bias of net input to these neurons than the bias to VIP neurons. All neurons were implemented as rate models with firing rate that was a rectified quadratic function of the summed input to the neuron,

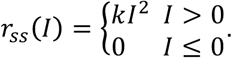

where *I* is the input strength, *r*_*ss*_ is the steady state firing rate, and *k* is a constant of proportionality. For ease of comparison with the SSN models developed by Rubin et al. (2015), we used *k* = 0.04 for all models.

For a given external input, the firing rates of all neurons in the network were obtained by evolving the network in time, with dynamics:

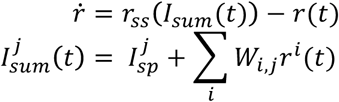

where *r*(*t*) is the time-dependent firing rate, 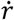 is the time derivate of the neuron’s firing rate, *r*_*ss*_ is the steady state firing rate that varies in time based on the inputs to the neuron, 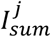 is the net input to neuron *j*, 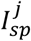 is a constant spontaneous input to neuron *j*, and *W*_*i,j*_ is the connection strength from presynaptic neuron *i* onto postsynaptic neuron *j*. To provide spontaneous activity to the network, and account for the higher spontaneous activity of VIP neurons^1^, we set 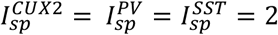 and 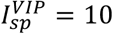. The network is evolved with Euler integration with updates of 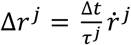 at each time step of Δ*t* = 0.1 *ms*, where the time constants of the different neuron types are 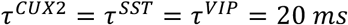 and 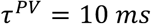.

